# *In vitro* cellular phenotypes of cortical neurons from R255X *MECP2* knock-in mice are improved by either expression of wildtype MeCP2 or ‘read-through’ with G418

**DOI:** 10.1101/2025.02.17.638700

**Authors:** Xin Xu, Jonathan Merritt, Steven J. Gray, Jeffrey Neul, Lucas Pozzo-Miller

## Abstract

Approximately 60% of individuals with Rett syndrome (RTT) carry a nonsense variant in the *MECP2* gene; thus, there is an unmet need to identify novel nonsense suppression compound(s) that can restore full length MeCP2 protein levels and function. Here, we characterized neuronal phenotypes in cultured cortical neurons from newborn knock-in mice harboring the *MECP2* R255X variant. After 2 weeks *in vitro*, R255X mutant neurons showed smaller cell bodies, shorter dendrites, fewer dendritic branches, and a lower density of excitatory synapses when compared to wildtype (WT) neurons. Transduction of AAV9-MeCP2-GFP in R255X mutant neurons made these cellular phenotypes similar to those in WT neurons, including soma size, dendritic length and branching, and excitatory synapse density. As proof of principle for the potential clinical use of ‘read-through’ compounds, cultured R255X mutant neurons treated with the aminoglycoside G418 for 72h *in vitro* showed cell body size and excitatory synapse density similar to WT neurons. We expect these combined approaches will identify effective compounds to suppress translation termination at a premature termination codon, which can be moved to further preclinical functional and behavioral studies in R255X *MECP2* knock-in mice.

**Summary Statement:** Expression of wildtype *MECP2* or treatment with G418 *in vitro* restored cell body size, dendritic length, and dendritic spine density in cortical neurons from R255X *MECP2* knock-in mice to levels comparable to wildtype neurons.

## INTRODUCTION

In-frame premature termination codons (PTCs), also known as nonsense pathogenic variants, comprise 11% of all disease-associated gene lesions (Mort et al., 2008). PTCs severely reduce gene expression via two mechanisms: 1) terminating translation of an mRNA before a full-length polypeptide is made; 2) destabilizing mRNAs due to activation of nonsense-mediated mRNA decay (NMD), a pathway that recognizes and degrades PTC-containing mRNAs (Keeling, 2016). It follows that translation termination and NMD represent two potential therapeutic targets for restoring deficient gene expression due to PTC.

Rett syndrome (RTT) is a neurodevelopmental disorder associated with intellectual disability primarily caused by loss-of-function variants in *MECP2*, the gene encoding the transcriptional regulator methyl-CpG-binding protein 2 (MeCP2) (Amir et al., 1999; Gold et al., 2024). Out of the reported ∼200 pathogenic variants in *MECP2*, only 8 account for the majority of RTT cases, with 60% of individuals carrying a nonsense variant, i.e. R168X, R255X, R270X, and R294X (Cuddapah et al., 2014). We hypothesize that suppressing translation termination at a PTC (alone or in conjunction with inhibiting NMD), will increase MeCP2 protein levels in cases of nonsense pathogenic variants to improve its proper function in RTT. Prior to testing novel nonsense suppression compound(s) in preclinical mouse models for RTT, we characterized cellular and synaptic phenotypes in cultured cortical neurons from newborn knock-in mice harboring the *MECP2* variant R255X (Pitcher et al., 2015).

## RESULTS AND DISCUSSION

We characterized cellular and synaptic phenotypes in cortical neurons from newborn R255X *MECP2* knock-in mice, which have significant phenotypic overlap with *Mecp2* null mice (Dong et al., 2020; Pitcher et al., 2015).

Cortical neurons in dissociated cultures were first transduced with AAV2-Camkii-GFP and fixed for morphological analyses after 2 weeks *in vitro* (DIV) (Fig. 1A). Cortical neurons from R255X mice (n=96 cells) have smaller cell bodies (WT: 384.1±8.1 μm^2^, R255X: 297.5±6.4 μm^2^, p<0.0001; Fig. 1B), shorter dendritic length (WT: 3,219.8±150.7 μm, R255X: 2,383.0±99.4 μm, p<0.0001; Fig. 2C), as well as fewer dendritic branches (WT: 29.9±1.4, R255X: 23.6±0.9, p=0.0003; Fig. 2D) compared to neurons from their WT littermates (n=104 cells). These observations are consistent with the smaller neuronal cell bodies described in cortical neurons from RTT individuals (Bauman et al., 1995a; Bauman et al., 1995b), male *Mecp2* knockout mice (Fukuda et al., 2005; Kishi and Macklis, 2004), and *Mecp2*-lacking neurons in female *Mecp2* heterozygous mice (Rietveld et al., 2015). Smaller dendritic trees with fewer branch points have also been observed in cortical neurons from RTT individuals (Armstrong et al., 1995), where they correlate with neuronal cell body size (Kaufmann et al., 1995), as well as in hippocampal neurons in dissociated cell cultures from newborn male *Mecp2* knockout mice (Larimore et al., 2009; Xu et al., 2017). These deficits are possibly caused by both a failure of the formation as well as of the maintenance of dendritic arbors in the absence of *Mecp2* (Baj et al., 2014).

**Figure 1.**
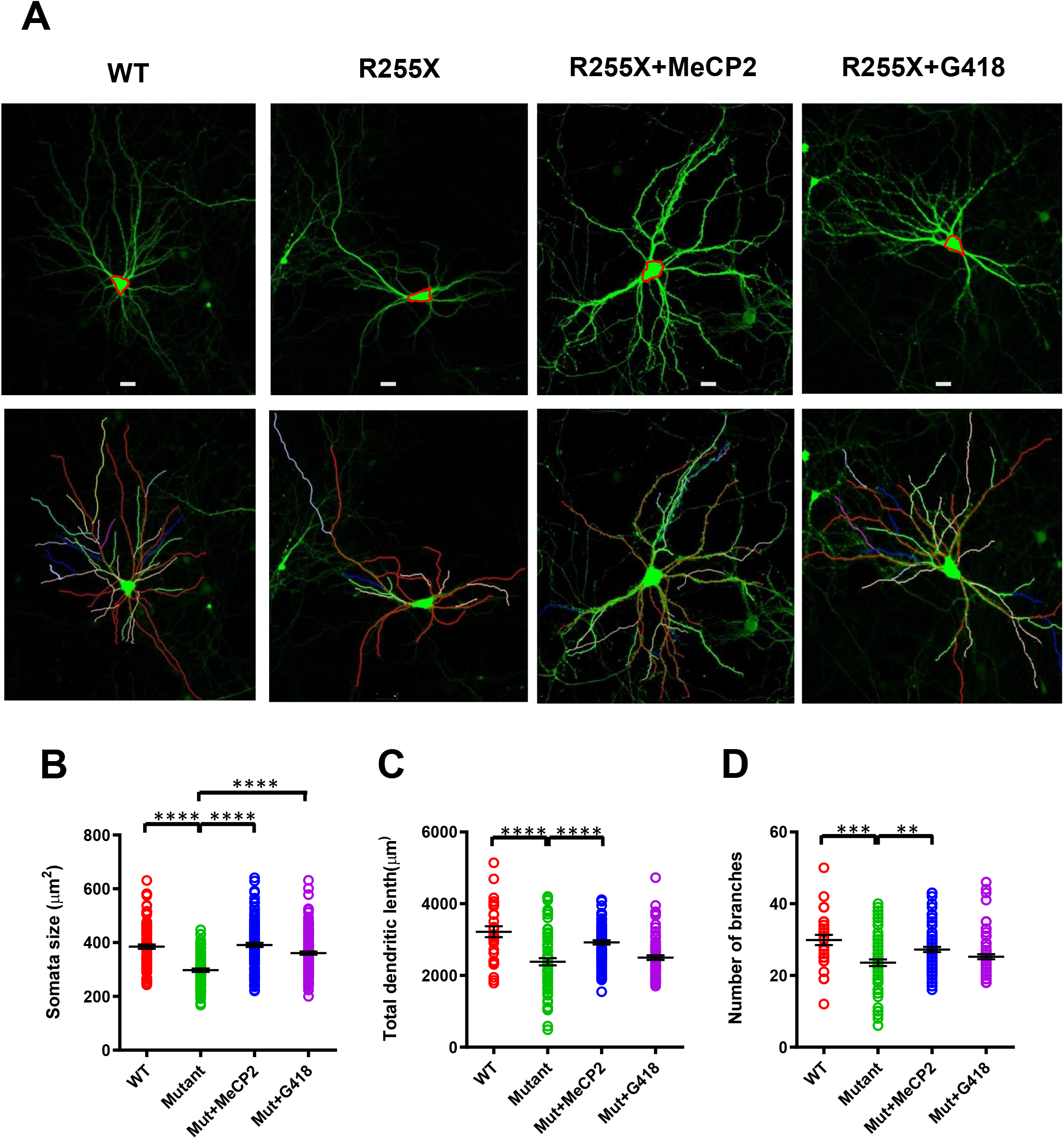
AAV9-mediated *MECP2* expression restores soma size, dendritic length, and dendritic branching in cortical neurons from R255X knock-in mice to those in WT levels, while readthrough with G418 restores only soma size in R255X neurons to WT levels. (A) Representative images show neurons transduced with AAV-CaMKII-GFP after immunostaining for GFP (green) in different conditions (top) and ImageJ dendrite tracing (bottom). (B) Soma size comparison in WT, R255X neurons, R255X neurons transduced with AAV9-MeCP2-GFP, and R255X neurons treated with G418. (C) Comparison of total dendritic length between genotypes and treatments. (D) Comparison of number of dendritic branches between genotypes and treatments. Scale bar= 20µm. ** P<0.01, *** P<0.001, **** P<0.0001

**Figure 2.**
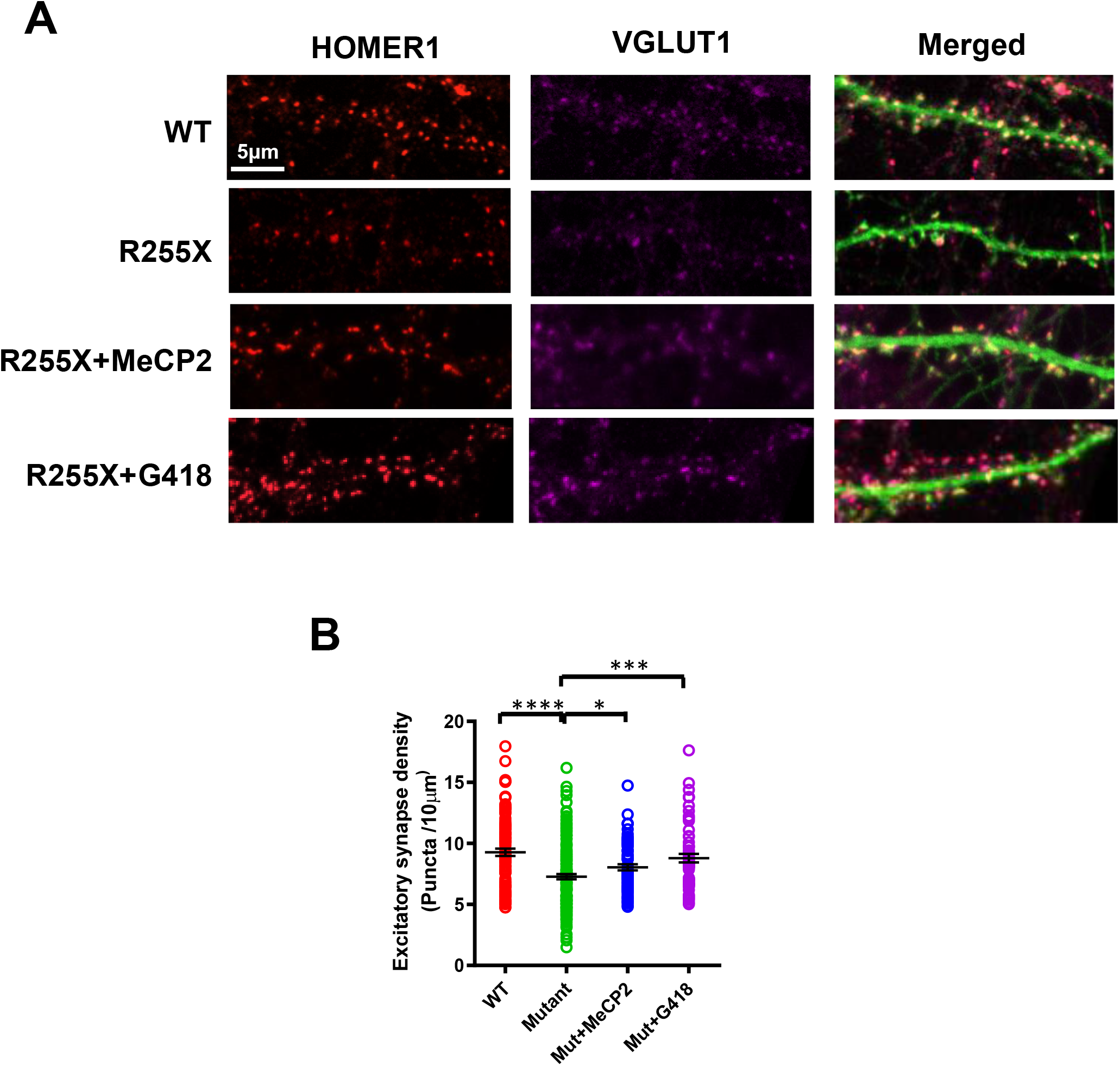
AAV9-mediated MeCP2 expression or readthrough with G418 restores the density of excitatory synapses in cortical neurons from R255X knock-in mice to those in WT levels. (A) Representative images of immunostaining for Homer1 (red) and VGLUT1 (purple) in GFP-expressing (green) neurons of different genotypes and treatments. (B) Comparison of numerical density of Homer1/VGLUT1 synaptic puncta between genotypes and treatments. * P<0.05, *** P<0.001, **** P<0.0001

The number of excitatory synapses, identified as VGLUT1-expressing presynaptic terminals opposed to dendritic spines of GFP-expressing neurons, and their surface levels of Homer1 were determined by triple color immunocytochemistry (Fig. 2A). Consistent with our previous reports (Xu et al., 2017; Xu and Pozzo-Miller, 2017) and other studies in hippocampal neurons from male *Mecp2* knockout mice *in vitro* (Baj et al., 2014; Chao et al., 2007), cortical neurons from R255X *MECP2* knock-in mice have fewer excitatory spine synapses compared to WT neurons (R255X: 7.28±0.21 VGLUT1/Homer1 puncta on GFP spines per 10 μm of dendrite; n=79 cells; vs. WT: 9.27±0.30 puncta/10 μm; n=135 cells; p<0.0001; Fig. 2B), reflecting delayed neuronal and synaptic maturation and/or excessive pruning of glutamatergic synapses.

Inducible re-expression of *Mecp2* from its endogenous promoter improves behavioral phenotypes and extends the lifespan of male *Mecp2* KO mice (Guy et al., 2007), suggesting that existing deficits can be reversed by *MECP2* gene therapy. Several groups have reported the delivery of different viral constructs encoding *MECP2* into different *Mecp2*-deficient mice (Collins et al., 2004; Gadalla et al., 2013; Garg et al., 2013; Luikenhuis et al., 2004; Luoni et al., 2020; Matagne et al., 2021; Matagne et al., 2017). We next confirmed that virally-driven expression of wildtype *MECP2* is sufficient to improve these cellular phenotypes in cortical neurons from R255X *MECP2* knock-in mice. Indeed, transduction of ssAAV9-CBh-GFP-WPRE-BGHpA::MeP229-hMeCP2-myc-spA (human *MECP2* under control of mouse *Mecp2* promoter, with GFP under control of the beta actin promoter) in cortical neurons from R255X *MECP2* knock-in mice (n=127 cells) made these cellular phenotypes similar to those of WT neurons, including soma size (391.2±7.6 μm2, p<0.0001; Fig. 1B), dendritic length (2,922.5±61.6 μm, p<0.0001; Fig. 1C) and branching (27.2±0.7, p=0.0016; Fig. 1D), as well as the density of excitatory synapses (n=61 cells; 8.04±0.24 puncta/10 μm; p=0.0200; Fig. 2B). These results provide additional support for the therapeutic potential of *MECP2* gene therapy for RTT (Gadalla et al., 2011; Sinnett et al., 2017).

An alternative strategy for reversing RTT pathologies is to overcome the effects of nonsense variants by readthrough of PTCs. G418 (geneticin), is a selective antibiotic of the aminoglycoside family, which allows ribosomal readthrough of PTCs during translation and can restore full-length functional protein (Howard et al., 2000). As proof-of-principle for the potential clinical use of readthrough compounds, cortical neurons from R255X *MECP2* knock-in mice that were treated with G418 (50ug/mL) for 48hs showed soma size (n=162 cells; 361.1±6.0 μm2, p<0.0001; Fig. 1B) and excitatory synapse density (n=106 cells; 8.52±0.22 puncta/10 μm; p=0.0001; Fig. 2B) comparable to WT neurons. However, the total dendritic length (2,541.6±48.6 μm, p=0.1546; Fig. 1C) and branching (25.2±0.6, p=0.1365; Fig. 1D) were not similarly rescued. Previous studies have reported that aminoglycosides (including G418) facilitate the synthesis of full-length MeCP2 in cells from R168X, R255X, and R294X knock-in mice (Brendel et al., 2011; Pitcher et al., 2015; Popescu et al., 2010). Our results further extend those observations by demonstrating that readthrough with G418 also rescues some of the neuronal phenotypes observed in cortical neurons from R255X *MECP2* knock-in mice. Finding effective compounds to suppress PTCs is an attractive strategy for RTT therapy because readthrough only occurs in the cells expressing the pathogenic variant allele, while the cells expressing the wildtype allele remain unaffected. However, the readthrough efficiency varies significantly between different compounds, which are also sensitive to the type of variant and the bases surrounding the premature termination codon.

In summary, our observations confirm that cortical neurons from R255X *MECP2* knock-in mice have cellular and synaptic phenotypes like those from *Mecp2*-deficient mice and RTT autopsy samples, which are amenable for screening novel more selective readthrough compounds. In addition, we confirmed that both AAV9-mediated *MECP2* expression and readthrough with G418 rescue some of these cellular phenotypes in cortical neurons from R255X *MECP2* knock-in mice to levels comparable to those from WT mice. Altogether, these results further support the rationale to search for novel and selective readthrough compounds to suppress translation termination at a PTC in neurons expressing nonsense pathogenic variants and improve RTT symptoms.

## MATERIALS AND METHODS

### Mice

All experimental mice were generated from *Mecp2*^R255X/X^ (Jackson Laboratory #012602, *Mecp2*^tm1.1Irsf/J^) mice that have been back-crossed to wildtype male C57BL6/J (Pitcher et al., 2015). Mice were handled and housed according to the Committee on Laboratory Animal Resources of the National Institutes of Health and had *ad libitum* access to food and water. All experimental protocols were reviewed annually and approved by the Institutional Animals Care and Use Committee of the University of Alabama at Birmingham.

### Neuronal cultures, virus transduction, and drug treatments

Cortical neurons were isolated from newborn (P0-1) female *MECP2* R255X heterozygous mice, male *MECP2* R255X mutant mice, and wildtype littermates, and cultured following standard methods. Cortex were rapidly dissected and dissociated in papain (20 U / ml) plus DNase I (Worthington Biochemical Corp., Lakewood, NJ, USA) for 25–30 min at 37°C, as described (Amaral and Pozzo-Miller, 2007; Xu et al., 2017; Xu and Pozzo-Miller, 2017). The tissue was then triturated to obtain a single-cell suspension, and the cells were plated at a density of 40,000 cells cm^−2^ on 18 mm poly-l-lysine/laminin coated glass coverslips, and immersed in Neurobasal medium (Life Technologies, Carlsbad, CA, USA) supplemented with 2% B27 (Life Technologies) and 0.5 mM glutamine (Life Technologies). Neurons were grown in 37°C, 5% CO2, 90% relative humidity incubators (Forma, Thermo Fisher Scientific, Waltham, MA, USA), with half of the fresh medium changed every 4–5 days. AAV2-Camkii-GFP (5×10e12 vg/mL; Addgene) transduction for morphological studies was performed at 7 days *in vitro* (DIV). Seven DIV cultures were transduced with ssAAV9-CBh-GFP-WPRE-BGHpA::MeP229-hMeCP2-myc-spA (human *MECP2* under control of mouse *Mecp2* promoter, with GFP under control of the beta actin promoter; 1.5×10e13 vg/mL) and fixed for immunocytochemistry 7 days later. Twelve DIV cortical cultures were exposed to G418 (50ug/mL; Fisher) for 48 hs and then fixed for immunocytochemistry.

### Immunocytochemistry

Cortical cultures were fixed with 4% (wt/vol) paraformaldehyde/sucrose for 10 min, incubated in 0.25% (v/v) Triton X-100 for 10 min, and then washed with phosphate buffered saline (PBS).

After blocking with 10% (v/v) goat serum in PBS, cells were incubated with primary antibodies against GFP (Abcam), Homer1 (Synaptic systems), and vesicular glutamate transporter (VGLUT1; Millipore), overnight at 4°C, rinsed in PBS, and incubated with secondary antibodies conjugated to Alexa 488, Alexa 594 and Alexa 647 (Jackson ImmunoResearch Laboratories, West Grove, PA, USA), respectively, for 2 h at room temperature. Coverslips were then mounted with Vectashield medium (Vector Laboratories, Burlingame, CA, USA) and imaged in a laser-scanning confocal microscope (Zeiss LSM-800, Carl Zeiss Microscopy, LLC, Thornwood, NY, USA) using a ×63 (1.4 NA) oil-immersion objective, as described (Xu et al., 2017; Xu and Pozzo-Miller, 2017).

### Image analysis

Cell body size was defined as the cross-sectional area of the outline around neuronal somata, using ImageJ. Dendrites were traced and measured using the *Simple Neurite Tracer* plug-in (ImageJ). The density of excitatory synapses was determined by the number of co-localized VGLUT1/Homer1 puncta per length of GFP-positive dendrite, using Imaris (Bitplane).

### Statistical analyses

Data are presented as mean ± standard error of the mean (SEM), and were compared using unpaired Student’s t-test for two groups or One Way ANOVA with Tukey post test for more than three groups. All analyses were performed using Prism software (GraphPad Software, San Diego, CA). P < 0.05 was considered significant.

## ACKNOWLEDGEMENTS

We thank Dr. Takafumi Inoue (Waseda University, Tokyo, Japan) for data acquisition and analysis software, and the *Civitan International Research Center* for the access to the confocal microscope. This work was supported by the *International Rett Syndrome Foundation* (LP-M).

